# Detour learning ability and the effect of novel sensory cues on learning in Australian bull ants, *Myrmecia midas*

**DOI:** 10.1101/2021.01.25.428158

**Authors:** Muzahid Islam, Sudhakar Deeti, Zakia Mahmudah, J. Frances Kamhi, Ken Cheng

**Affiliations:** Department of Biological Sciences, Macquarie University, Sydney, Australia; Neuroscience Department, Oberlin College, Oberlin, Ohio, USA

**Keywords:** Detour Learning, Successful Foragers, Path Straightness, Motor Routine

## Abstract

Many animals navigate in a structurally complex environment which requires them to detour around physical barriers that they encounter. While many studies in animal cognition suggest that they are able to adeptly avoid obstacles, it is unclear whether a new route is learned to navigate around these barriers and, if so, what sensory information may be used to do so. We investigated detour learning ability in the Australian bull ant, *Myrmecia midas*, which primarily uses visual landmarks to navigate. We first placed a barrier on the ants’ natural path of their foraging tree. Initially, 46% of foragers were unsuccessful in detouring the obstacle. In subsequent trips, the ants became more successful and established a new route. We observed up to eight successful foraging trips detouring around the barrier. When we subsequently changed the position of the barrier, made a new gap in the middle of the obstacle, or removed the barrier altogether, ants mostly maintained their learned motor routine, detouring with a similar path as before, suggesting that foragers were not relying on barrier cues and therefore learned a new route around the obstacle. In additional trials, when foragers encountered new olfactory or tactile cues, or the visual environment was blocked, their navigation was profoundly disrupted. These results suggest that changing sensory information, even in modalities that foragers do not usually need for navigation, drastically affects the foragers’ ability to successful navigate.

**Subject Category:** Neuroscience and Cognition

## Introduction

In detour learning, a significant paradigm in animal cognition, a direct route is blocked, and a new route must be taken to reach the goal. In a natural environment, animals face various situations where obstacles block their regular shortest path to reach a destination, and in most cases, they can negotiate these obstacles (Zucca et al., 2005; Kabadayi et al., 2018). Several ecological and contextual factors might affect the efficiency of animal detour learning, such as the nature of the barrier, goal distances, detour experiences, reward visibility, and motivation (Kabadayi et al., 2018). To meet these challenges, animals may integrate both external and internal cues that help to redirect their movement (Giurfa, 2015; Kabadayi et al., 2018) but it is unknown what cues are used for establishing a new route.

Two forms of detour learning have been identified: spatial learning, characterized by gradual learning, and spatial reasoning, characterized by quick learning. Spatial learning occurs if an individual develops a tendency to select the shortest route over multiple trips. In contrast, spatial reasoning occurs if an individual selects the shortest path to a destination spontaneously on its first trip based on memory and visual inspection (Kabadayi et al., 2018). Detour abilities have been studied in a wide range of taxa including vertebrates such as apes (Kohler, 1925), rats and mice (Juszczak et al., 2016), dogs (Pongracz et al., 2001; Siniscalchi et al., 2013) goats (Naworth et al., 2016; Langbein et al., 2018), horses (Baragli et al., 2011; Baragli et al. 2017), marsupials (Wynne and Leguet, 2004), fish (Bisazza et al., 1997; Sovrano et al., 2018; Lucon-Xiccato et al., 2019), birds (Vallortigara et al., 1999; Zucca et al., 2005) and invertebrates such as spiders (Tarsitano and Jackson, 1997; Aguilar-Argüello et al., 2019), and bees (Kevan, 2015; Ong et al., 2017). In most detour experiments, animals have been tested repeatedly to examine their learning abilities (Kabadayi et al., 2018). Several studies have shown gradual improvements in detouring over multiple trials (Parker et al., 2005; Smith and Litchfield, 2010; Boogert et al., 2011). For example, horses reach their goal faster over trials when they encounter an obstacle (Baragli et al., 2011). On the very first trip, some horses were unsuccessful in detouring the barrier but from the second trip they improved their success. Similarly, dingos in a detour learning experiment improved their performance significantly over four trials (Smith and Litchfield, 2010). Another detour learning experiment was done on mice, which exhibited a preference for one or the other side of the barrier and optimized the direction of movement based on their previous experiences, showing gradual learning over trials (Juszczak et al., 2016). Another study investigated the ability to progress around a barrier to a desired food source in four quokkas, *Setonix brachyurus*, in their natural environment (Wynne and Leguet, 2004). Among them, three quokkas preferred to detour either right or left around the obstacle, showing spatial learning, while the fourth showed no behavioural laterality but exhibited spatial reasoning.

Ants are highly efficient navigators in complex natural environments (Freas and Cheng, 2019; Lihoreau and Schwarz, 2020; Buehlmann et al., 2020). There have been few studies of detour ability in ants. A detour strategy was observed in the crazy ant, *Paratrechina longicornis* in obstacle navigation during cooperative transport (McCreery et al., 2016). These ants proved effective at implementing a flexible or stochastic strategy that works for both simple and complex barriers. Another detour experiment on *Cataglyphis fortis* ants showed that when a barrier was set between the nest and releasing point, homing ants, tested just once, usually walked towards the barrier, and when coming close to the barrier, deviated around one of the sides and moved forward to the goal (Schmidt et al., 1992). Desert ants (*Melophorus bagoti* and *Cataglyphis fortis*) can also learn a route detour around a pit trap (Wystrach et al., 2020).

The Australian nocturnal bull ant, *Myrmecia midas*, is an excellent navigator that forages on eucalyptus trees by using primarily visual information (Freas et al., 2017a; Freas et al., 2018; Freas and Cheng, 2019; Islam et al., 2020). Nest entrances of these bull ants are usually located close to a eucalyptus tree, called their nest-tree (Freas et al., 2018). Ants forage individually in the tree canopy of either their nest tree or other nearby trees for insect prey and sap. There is no evidence of trail pheromone for recruitment or any other recruitment interaction among the foragers. Ants from each nest typically travel to a foraging tree in evening twilight and back to their nest before morning twilight (Freas et al., 2017a; Freas et al., 2018).

This study investigated the ability of *M. midas* to detour around a barrier during the ascending navigation on their foraging tree, with a focus on four questions. First, would the ants successfully move around the detour on each trial? Second, would the ants show spatial learning or spatial reasoning? Third, how would the bull ants modify their route when the barrier is removed partially or completely? Fourth, how would ants behave when encountering visual and non-visual sensory changes along their new route? We hypothesized that ants would learn to overcome the barrier gradually with repeated trials, as seen in previous detour learning studies (e.g., Parker et al., 2005; Smith and Litchfield, 2010; Boogert et al., 2011). Given their reliance on visual cues for navigation, we predict that the ants would follow their newly learned route when encountering any changes of barrier position or any olfactory or tactile changes on their foraging route, but that they would navigate less successfully when visual cues around the tree were blocked.

## MATERIAL AND METHODS

### Animal and experimental site

The experimental site was located at Macquarie University campus, Sydney, Australia (33°4611’’ S,151°06’40’’ E). Nocturnal bull ants, *Myrmecia midas*, typically build their nest close to a Eucalyptus tree. We selected two nests 200 m apart which were located close to a smooth-skinned eucalyptus tree, *E. virminalis.* In Experiment-1, we conducted experiments on both Nest-A and Nest-B during 2018–2019 and on only Nest-A during 2019–2020. In Experiment-2, we conducted experiments on Nest-A during 2019–2020. A tree is usually located less than 30 cm from nest entrances of *M. midas* colonies (Freas et al., 2017a; Freas et al., 2017b). This tree is termed the “nest-tree.” Ants typically emerge from the nest entrance, walk a few centimetres on the ground, and then many foragers climb vertically up the nest-tree. We call this behaviour “ascending navigation,” which is perpendicular to the direction of navigation on the ground. To record the path of individual foragers during their ascending foraging trips, a grid was drawn from the bottom of the tree to 1.5 m high on the tree trunks of the nest-trees of both Nest-A and Nest-B. Each grid consisted of 10-cm squares (Fig. 1) that covered the diameter of the tree trunk with code numbers on the grid to identify locations on the grid. The circumference of the Nest-A and Nest-B nest-tree trunks were 200 cm and 160 cm, respectively.

**Fig. 1.**
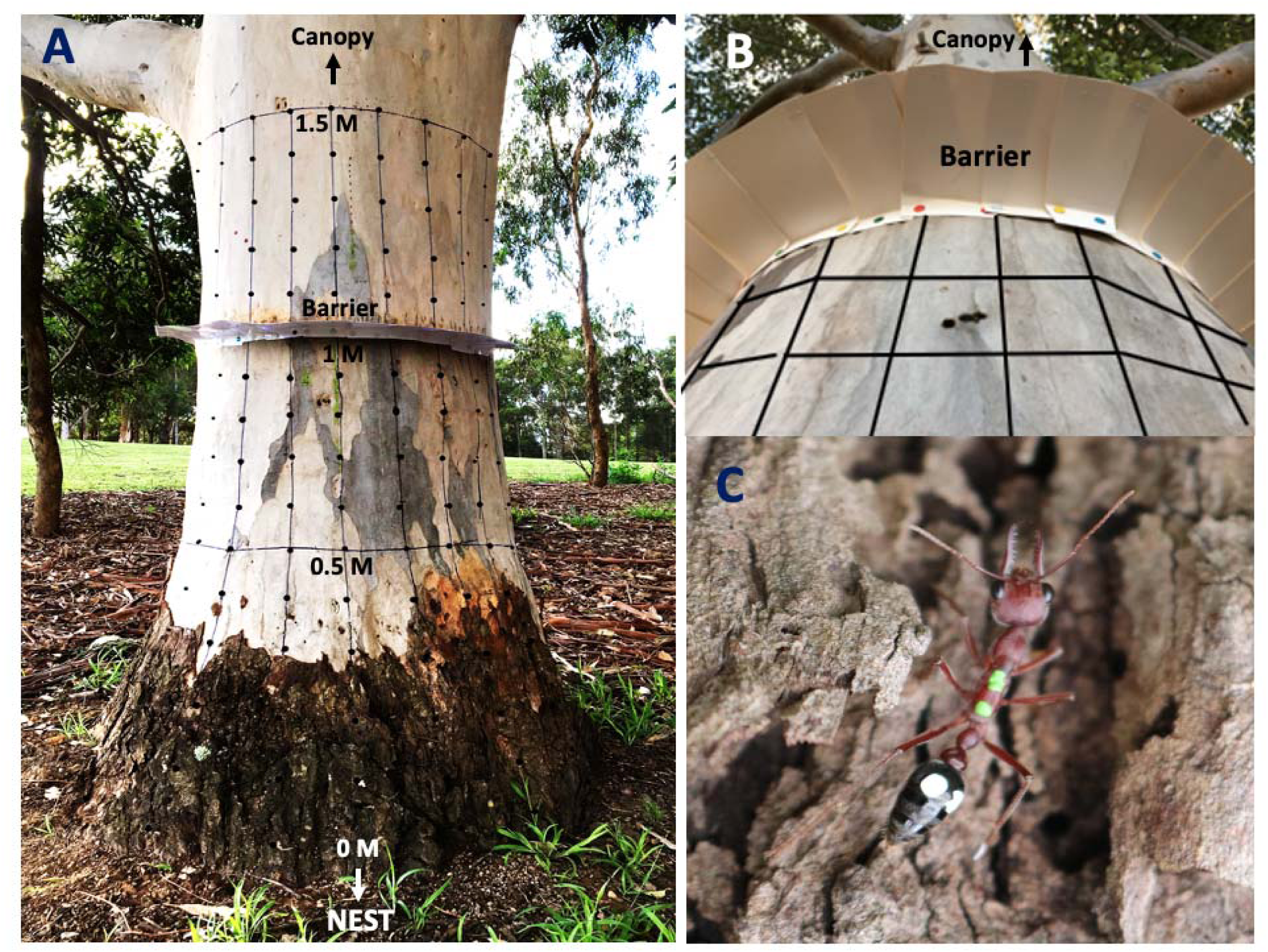
The experimental set up for the detour learning experiments on Australian nocturnal bull ants, *Myrmecia midas*, on their nest-tree: A. The tree trunk of a nest-tree (Nest-A) with a grid from the base of the tree (0 m) to 1.5 m. A white cardboard barrier was placed perpendicular to the tree trunk at 1 m from the base of the tree trunk which covered 180 degrees at the nest-side. B. A close view of the barrier that the foragers were confronted with. C. A uniquely colour-coded nocturnal bull ant *Myrmecia midas* ascending towards the top of the tree during detour learning. The photo was taken when the worker ant started at the bottom of the foraging tree in the evening twilight.

### Experimental procedure

#### Experiment 1

In the Control Condition, we observed individually marked ants between 7:00 p.m. and 9:30 p.m. for seven consecutive nights to record the frequency and paths of natural, unobstructed foraging routes on the nest-tree. On the very first night, we captured the ants from the nest-tree trunk by using plastic tubes and put them into an icebox with ice for 3–5 minutes to cool. Then foragers were painted with a unique colour code on the dorsal side of their body with Citadel Paint (Games Workshop, UK). For detour learning tests, we chose those marked ants that ascended to forage on the nest-tree at least three times out of seven consecutive nights (104 foragers out of 135, 18 ants from Nest-A 2018–2019, 14 ants from Nest-B and 72 ants Nest-A 2019–2020). In the Learning Condition, a piece of white art paper (15 cm in diameter and 0.25 mm thick) was placed at 1 m from the bottom of the tree trunk which covered 50% of the perimeter of the nest-tree trunk, obstructing their natural foraging corridor on the nest side (Fig. 1). We tested the selected 104 foragers in up to eight trials each with a detour barrier over 21 consecutive nights immediately after the Control Condition for each nest.

#### Experiment 2

After the Control and Learning Conditions on Nest-A in 2019–2020, we tested foragers one time in each of six different conditions in order: Extension Condition (we moved the barrier from 1 m to 1.5 m from the bottom of the tree trunk, n=38), Gap in Middle Condition (we made a 20-cm gap in the middle of the barrier, n= 40), No Barrier Condition (we took off the entire barrier, n=36), Tactile Condition [we added a cream-coloured cloth (100 cm × 120 cm) which covered between 0.5 m and 1.5 m height on the front side of the tree trunk, n=38], Olfactory Condition (we sprayed perfume on two different places of the tree trunk at 50 and 80 cm height which made two parallel lines and covered 50% of nest side, n=25), and Visual Condition (we installed a 2-m wide and 1.25-m tall black screen on both the right and left sides symmetrically on the ground and 50 cm from the edge of the tree trunk on both sides, n=36). The barrier was not removed in the Tactile, Olfactory and Visual Conditions. After testing in each Condition, we allowed the foragers 10 days of relearning the route with the barrier (Table S1, Fig. S1). During relearning, we kept the barrier at 1 m from the bottom of the tree trunk as in the Learning Condition. We used the individually marked foragers observed in the Control and Learning Conditions for all of the additional Conditions. Not all ants were tested in each condition because some ants did not forage every night of the experimental conditions. Full details of methods are provided in the supplementary materials.

### Statistical Analysis

Graph-Click (www.arizona-software.ch/graph-click) was used to digitize the path of individual foragers. By using a custom-written MATLAB (MATLAB 2019b) program, we plotted the paths of the foragers and calculated the path straightness of individual ants in both the Control Condition and the Learning Condition in the two different nest-trees separately. We determined the path straightness for the path from the bottom of the tree to one-meter height as the ratio of the straight-line distance to the sum of all segments of the path (Palavalli-Nettimi et al., 2018) transformed so that 0.5 is the maximum value. For the Control Condition, the straight-line distance for individual foragers was ∼1 m, but in the Learning Condition, the straight-line distances were 112 cm and 108 cm (from the bottom of the tree trunk to edge of the barrier) in Nest-Tree A and Nest-Tree B, respectively. For all of the other conditions except the Extension Condition the straight-line distance was also 112 cm (straightness measured up to 1 m). For the Extension Condition this distance was 158 cm (straightness measured up to 1.5 m). We considered ants as successful on a trip if they passed the barrier located at 1 m from the bottom of the tree trunk.

For the Learning Condition, we conducted repeated-measures ANOVAs in SPSS to compare the mean path straightness across the 3^rd^ Control run in the Control Condition and 5 successful foraging trips (1^st^, 2^nd^, 3^rd^, 5^th^ and 8^th^) of the Learning Condition in Nest-A 2018– 2019, Nest-A 2019–2020 and Nest B. We determined whether there was a significant difference in path straightness of each successful learning trip in the Learning Condition compared to the Control Condition by using a Bonferroni correction. The foraging duration was timed when individual ants started from the bottom (0 cm) of the nest tree until they reached 1.5 m from the bottom in both the Control Condition and the Learning Condition. We conducted a one-way repeated-measures ANOVA with a Bonferroni correction to compare durations of the 3^rd^ trip of the Control Condition and 5 different successful foraging trips (1^st^, 2^nd^, 3^rd^, 5^th^ and 8^th^) during the Learning Condition in Nest-A 2018–2019, Nest-A 2019–2020 and Nest B. All of the foragers detoured either on the left or right side of the barrier during their learning trips. We reflected the paths of the ants going to the right so that all ants, conceptually, headed to the left side of the barrier. This practice allowed data to be pooled to examine how the foragers adjusted their upward path with the barrier over trips. To compare changes in paths over foraging trips (3^rd^ control trip, 1^st^, 2^nd^, 3^rd^ 5^th^ and 8^th^ successful foraging trips), the mean positions on the *x*-axis (horizontal position) of the ants on the tree trunk at 0 cm, 10 cm, 20 cm, 30 cm, 40 cm, 50 cm, 60 cm, 70 cm, 80 cm and 90 cm, we used a two-way repeated-measures ANOVA with height and trip as factors. For all repeated measure ANOVA, while the p-value represents whether the difference between conditions is statistically significant or not, partial eta squared (ηp^2^) illustrates about the magnitude of this difference, or effect size. A value of ηp^2^ ≥0.14 is considered a large effect.

In Experiment-2, to compare path straightness and duration, paired-samples t-tests were conducted to compare the differences between last learning trips or relearning trips and test-condition trips (for instance, Middle Gap tests vs. immediately preceding relearning trip). We report effect sizes (Cohen’s d) along with statistical significance levels. Data were analyzed using SPSS Statistics (Version 26), MATLAB (2019b) and R-Studio (Version 1.2.1335/2019).

## RESULTS

### A. Experiment-1: Detour Learning

#### Successful versus unsuccessful foraging trips

Pooling results from both Nest-trees A and B in both experiments, first foraging trips with a barrier were challenging for individual foraging ants, with 46% unsuccessful (Fig. 2). The ants improved steadily over trips, with almost all successful by the 5^th^ trip.

**Fig. 2.**
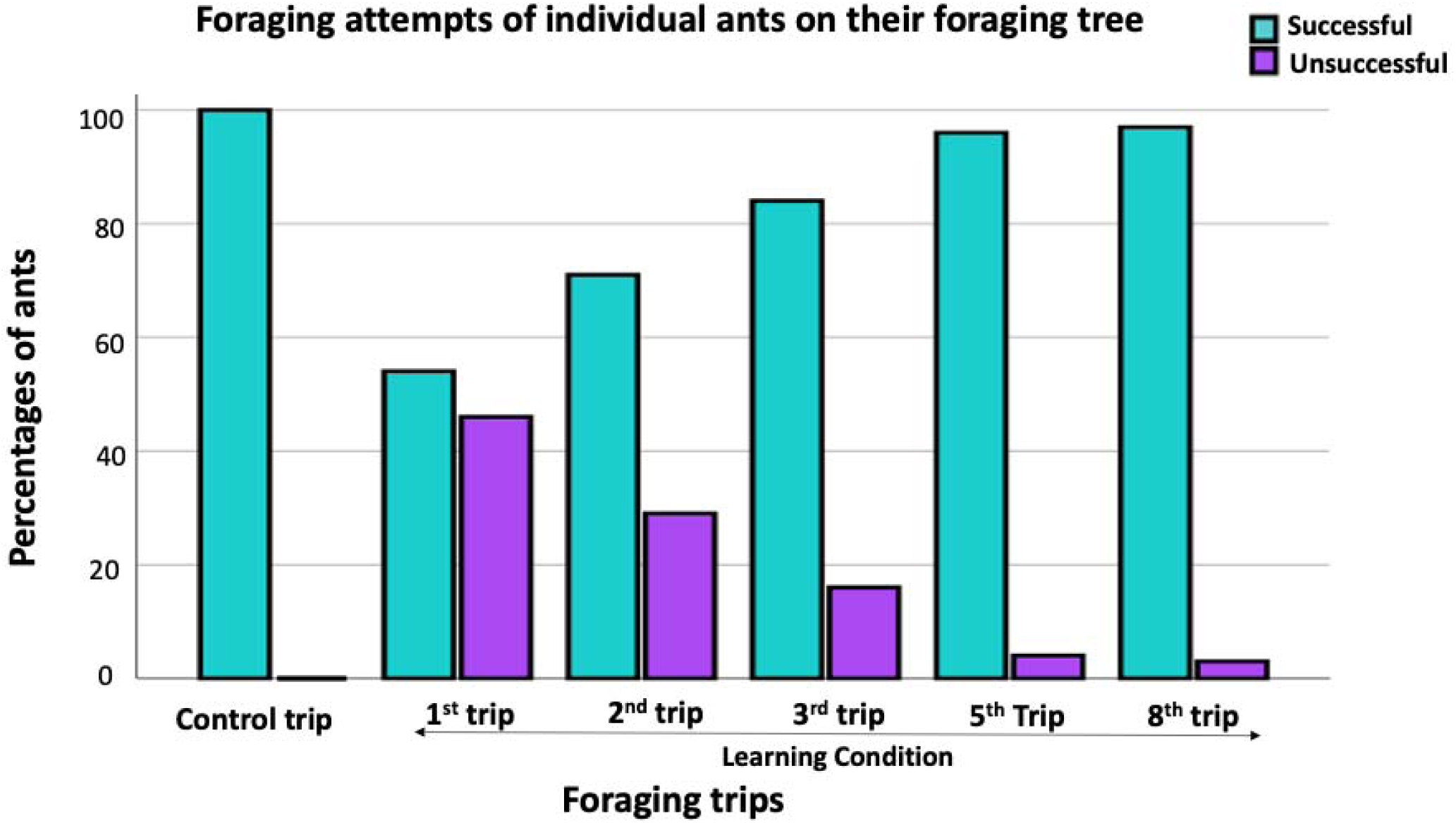
The percentages of successful and unsuccessful foraging ants (N=104, Nest-Tree A, 2018–2019=18, Nest-Tree B=14, Nest-Tree A, 2019–2020=72) in the 1st, 2nd, 3rd, 5th and 8th attempts of foraging trips.

#### Position of foragers on the tree trunk on successful foraging trips

In the Control Condition, foragers’ position was close to 0 on the *x*-axis from 0 cm to 90 cm of the tree trunk (data not shown). Over trips in the Learning Condition, foragers’ average heading position became farther from 0 on the x-axis, which indicates ants learning to detour. A two-way repeated-measures ANOVA examined the effects of trips and height on the *y*-axis on the dependent variable of *x*-axis position. In Nest-Tree A 2018–2019, there was a statistically significant interaction between trips and *y*-axis height [F (36,600) =6.273, p < .05, *ηp^2^* = 0.273]. There were also significant main effects of trips [F (4,600) =149.03, p < .05, *ηp^2^* = 0.499] and height on the *y*-axis [F (9,600) =11.495, p < .05, *ηp^2^* = 0.407]. In Nest-Tree A 2019–2020, a statistically significant interaction was also observed between trips and position on the *y*-axis [F (36,3599) =9.85, p<.05, *ηp^2^* = 0.191]. Significant main effects were also found for trips [F (4,3599) =400.78, p < .05, *ηp^2^* = 0.308] and position on the *y*-axis [F (9,3599) =117.71, p < .05, *ηp^2^* = 0.227]. Like Nest-A, there was a statistically significant interaction between trips and *y*-axis position of ants in Nest-Tree B [F (4,320) =5.743, p < .05, *ηp^2^* = 0.392] and also significant main effects of trips [F (4,320) =135.28, p < .05, *ηp^2^* = 0.628] and height on the tree [F (9,320) =13.09, p < .05, *ηp^2^* = 0.594]. In all experiments, the position of foragers on the tree trunk was not significantly different (p > 0.05) between the Control Condition and the 1st successful foraging trip. However, the 2nd, 3rd, 5th and 8th successful foraging trips were significantly different (p < 0.05) compared to the Control Condition. Importantly, in the Learning Condition, there were no significant differences of ants’ mean position on the *x*-axis of the tree trunk between the 5th and the 8th successful foraging trips (p > 0.05). The head positions of foragers on the *x*-axis were not significantly different across trips at 0 cm and 10 cm height, but from 20 to 90 cm height, foragers’ *x*-axis positions were significantly different (p < 0.05) across trips, suggesting that detouring started between 10 cm and 20 cm height on the tree trunk (Fig. 3).

**Fig. 3.**
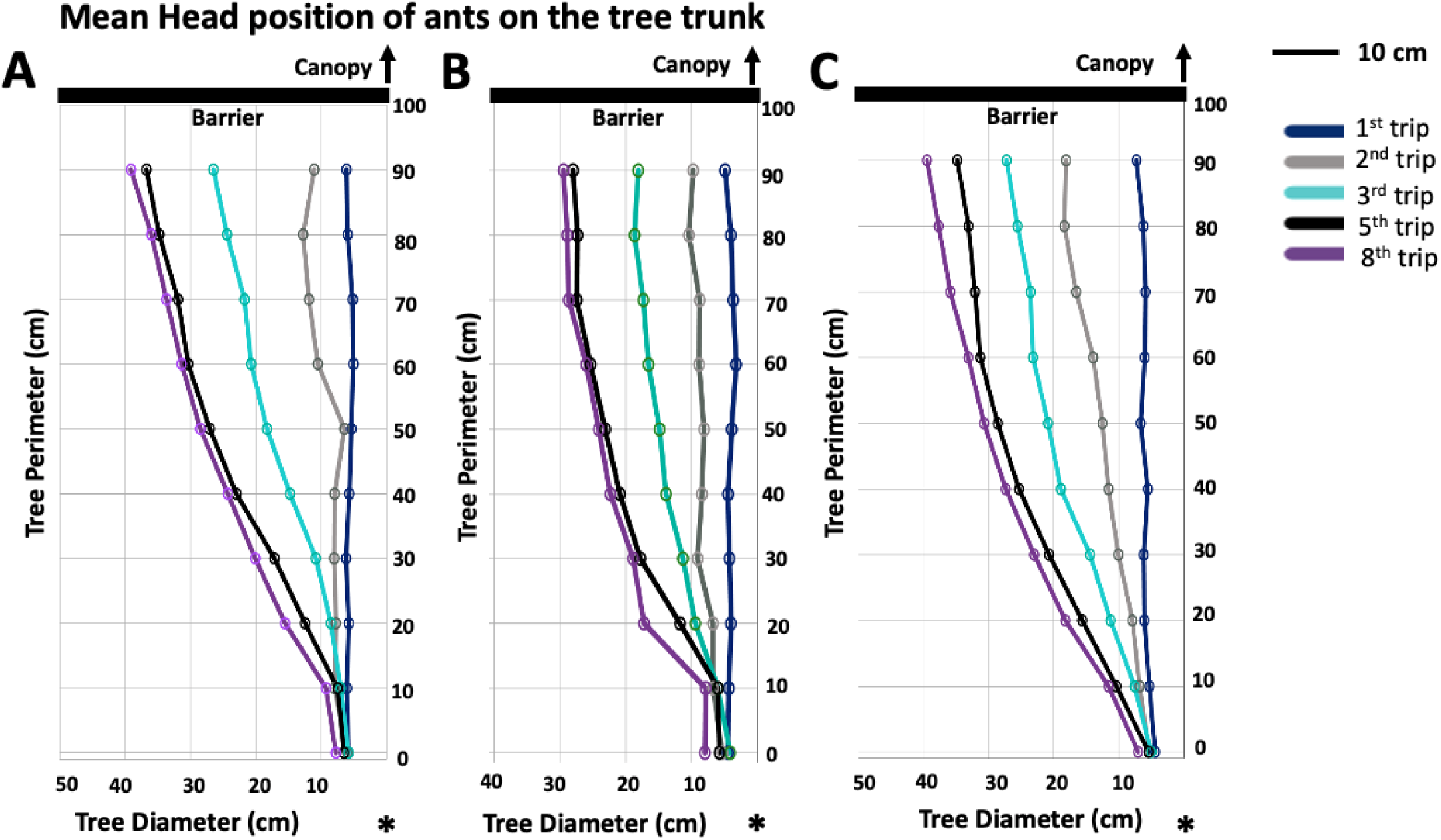
Interaction between trip number in the Learning Condition and height on the tree trunk in the *x*-axis position of ants on the nest-tree trunk in Nest-tree A 2018–2019 (A), Nest-tree B (B) and Nest-tree A 2019-2020 (C). Nest-tree A trunk was 200 cm in circumference where Nest-tree B was 160 cm in circumference and the barrier covered 50% on the front side at 1 m from the bottom of the tree trunk (see details in Experimental Procedure). The figure represents one half of the front (nest) side of the tree trunk. The *x*-axis represents the horizontal position on the tree trunk and the *y*-axis represents the height on the tree trunk. We reflected the paths of the ants going to the right so that all ants are, conceptually, headed to the left side of the barrier. The coloured dotted lines illustrate the mean of *x*-axis positions of ants on the tree trunk in different successful learning trips. The black line indicates the position of the barrier on the tree trunk at 1 m. Ant paths are shown from 0 cm to 90 cm of the tree trunk. The arrow indicates the canopy direction and the ‘*’ indicates the nest position.

The mean data in figure 3 hide notable individual differences in detour learning. Foragers showed 3 types of detour learning: A. Gradual learning, a forager that learned the detour path gradually (figure S2A); B. Quick learning, a forager that adjusted after 2 trips (figure S2B); C. Barrier-avoiding learning, a forager that followed the foraging corridor up to the barrier and then detoured to one side (Fig. S2C).

#### Path Straightness

The path straightness of foragers in detour learning was compared to their performance in the control condition. Foragers ascended directly to the canopy on the tree-trunk in the Control Condition, thus having a path straightness near the maximum value of 0.5 (figure 4). Path straightness dropped sharply in the 1^st^ successful foraging trip in the Learning Condition, and then improved over trips. In the 5^th^ and 8^th^ successful foraging trips foragers performed straighter paths like those in the Control Condition. A one-way repeated-measures ANOVA showed a statistically significant difference between the Control Condition and successful foraging trips in the Learning Condition of foragers in all experiments: Nest-Tree A 2018– 2019 (F(5,75) = 151.05, p < 0.001, ηρ^2^ = 0.988), Nest-Tree B (F(5,40) = 38.56, p < 0.001, ηρ^2^ = 0.903) and Nest-tree A 2019–2020 (F (5,355) = 334.06, p < 0.001, ηρ^2^ = 0.825). Post hoc comparisons using Bonferroni correction showed that the first two successful trips in the Learning Condition were significantly less straight when compared with the Control condition in all experiments, while the third trip was significantly less straight in the 2018-19 field season (Fig. 4). The 5^th^ and 8^th^ trips did not differ significantly from the Control Condition in either colony or season.

**Fig. 4.**
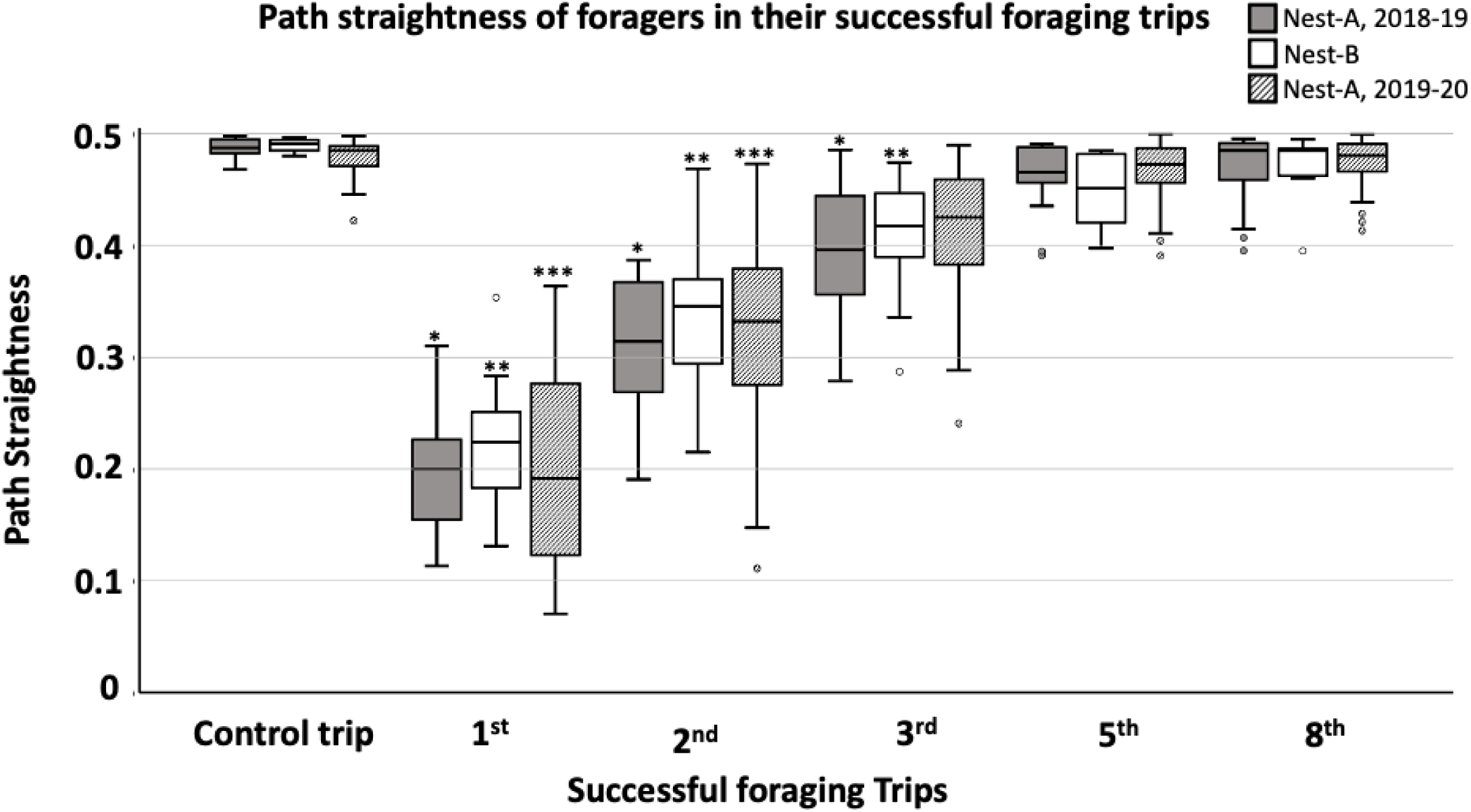
Path straightness of *Myrmecia midas* during their Control and Learning Conditions. The ‘*’ indicates that the path straightness of the 1^st^, 2^nd^ and 3^rd^ successful foraging trips were significantly different from the Control Condition in Nest-A 2018–2019 and the ‘**’ indicates a significant difference in path straightness compared to the Control Condition in Nest-B. Similarly, ‘***’ represents the significant differences of path straightness in their 1^st^, 2^nd^ and 3^rd^ learning trips compared to the Control Condition in Nest-A 2019–2020. The box plots indicate medians (solid black line), box margins (25th and 75th percentiles) and whiskers (5th and 95th percentiles) in this and all following figures.

#### Duration of foraging trips

In the 1^st^ successful foraging trip ants took 2–3 fold more time compared to the Control Condition (figure 5). Over successful learning trials foragers took significantly less time to pass the barrier compared to the Control Condition in all experiments: Nest-tree A 2018– 2019 (F (5,75) = 84.02, p < 0.001, *ηp^2^*= 0.851), Nest-Tree B (F (5,40) = 34.012, p < 0.001, *ηp^2^* = 0.812), Nest-tree A 2019–2020 (F (5,355) = 154.65, p < 0.001, *ηρ^2^*= 0.685). Post hoc comparisons using Bonferroni correction indicated significant differences from the Control Condition in the first two trips in the Learning Condition in all experiments, while the third trip differed significantly from the Control Condition in two of the three experiments (Fig. 5). The 5^th^ and 8^th^ trips did not differ significantly from the Control Condition in any experiment.

**Fig. 5.**
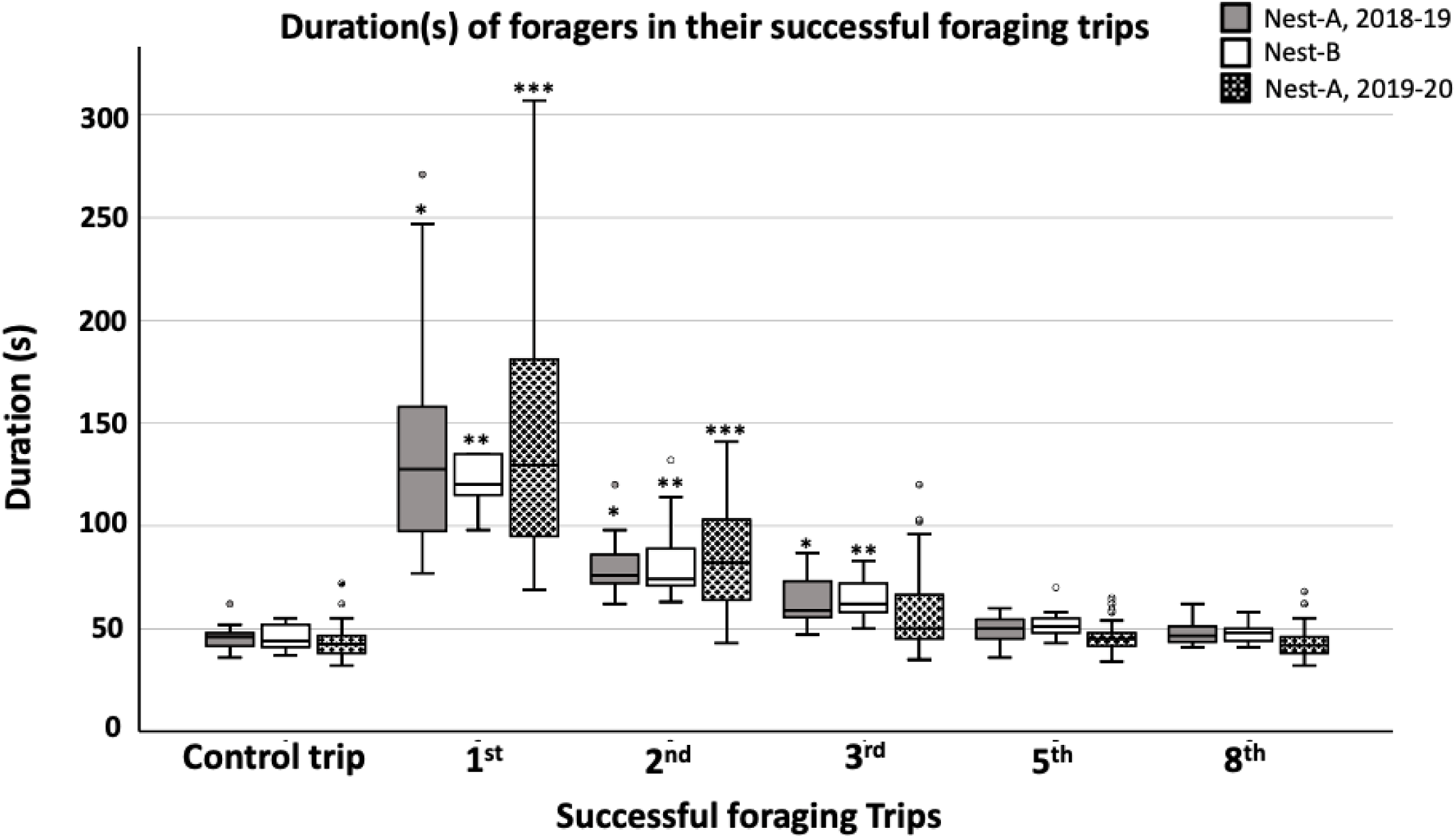
The duration (up to 1.5 m height) on the ants’ successful foraging trips in both Nest-A and Nest-B. The ‘*’ indicates that the duration of 1^st^, 2^nd^ and 3^rd^ successful foraging trips were significantly different from the Control Condition in Nest-A 2018–2019 and the ‘**’ indicated the significant differences in duration compared to the Control Condition in Nest-B. Similarly, ‘***’ represents the significant differences of duration in their 1^st^ and 2^nd^ learning trips compared to the Control Condition in Nest-A 2019–2020.

#### Laterality sequence in detour learning

Foragers performed with individual side preferences during their successful learning trips. Only two out of 104 foragers modified their foraging direction and detoured both to the right and to the left sides. In both colonies and seasons, similar numbers detoured on the left and right sides. A binomial test showed the proportions of right and left detours do not differ significantly from the chance of expectation of 50:50 in any group (Table S2).

### B. Experiment-2: Effect of Changes on Learning

#### Effects of barrier position on detour learning

Changing the barrier position (Extension Condition), adding a gap to the barrier (Middle Gap Condition), and removing the barrier (No Barrier Condition) did not greatly affect the paths of the ants. The ants took a similar path as in the previous experiment (Fig. 6A-C). Even with no barrier at all, the ants headed to the position of the edge of the barrier. Although path straightness (Fig. 7) and trip duration (Fig. 8) in the first test (Extension Condition) differed from the previous learning trip (path straightness, paired t test, t (35) = 5.838, p < 0.001; duration, paired t test, t (35) = –3.468, p = 0.001), these differences disappeared in the subsequent two tests, including the test with the drastic change of not having any barrier at all.

**Fig. 6.**
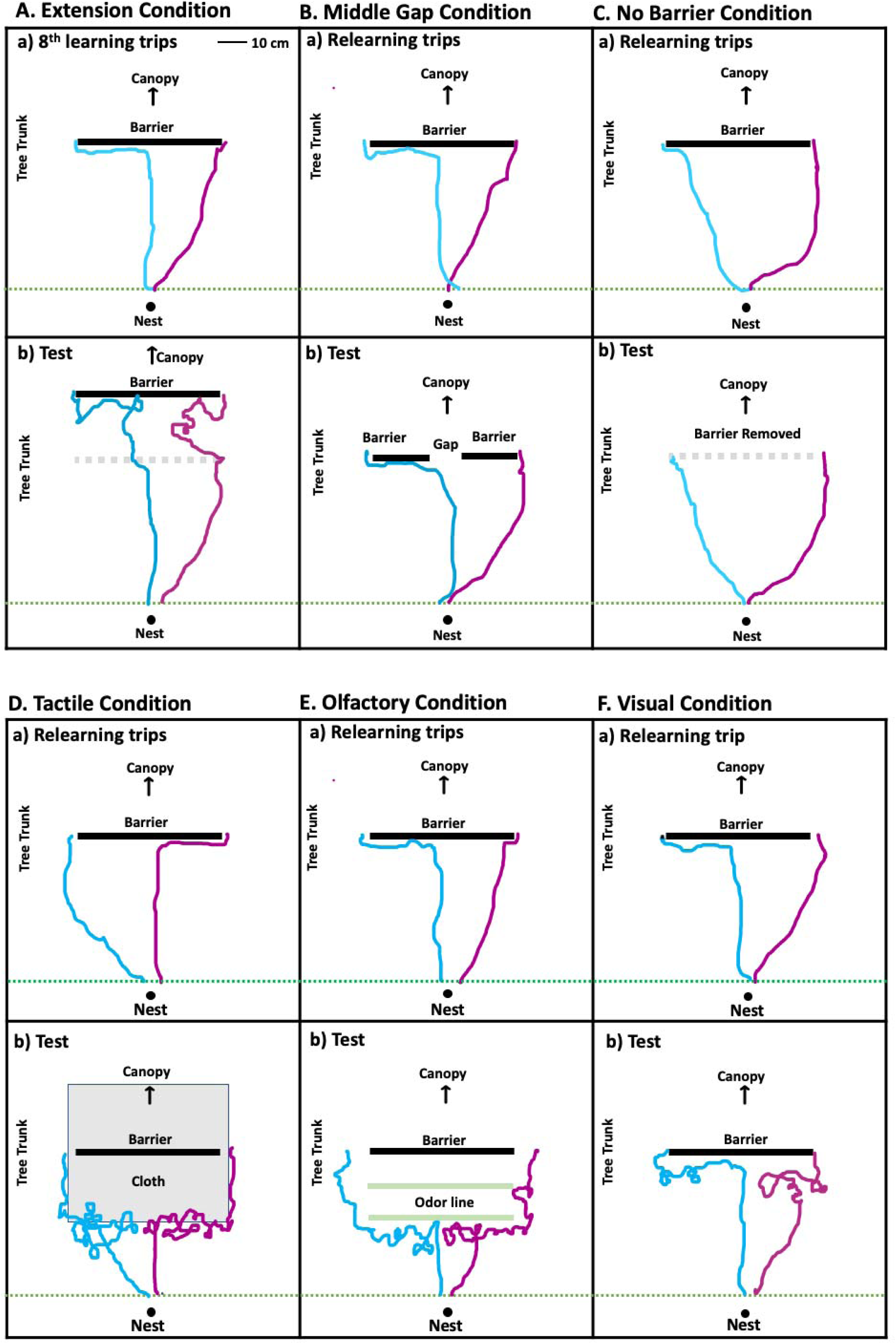
Exemplary paths of foragers in 6 different conditions with manipulations: (A) Extension Condition, (B) Middle Gap Condition, (C) No Barrier Condition, (D) Tactile Condition, (E) Olfactory Condition, and (F) Visual Condition. In each condition, (a) represents foragers’ habitual path after 8^th^ learning trips or relearning trips in the presence of barrier at 1m whereas (b) represents the ant’s response to the particular Changes.

**Fig. 7.**
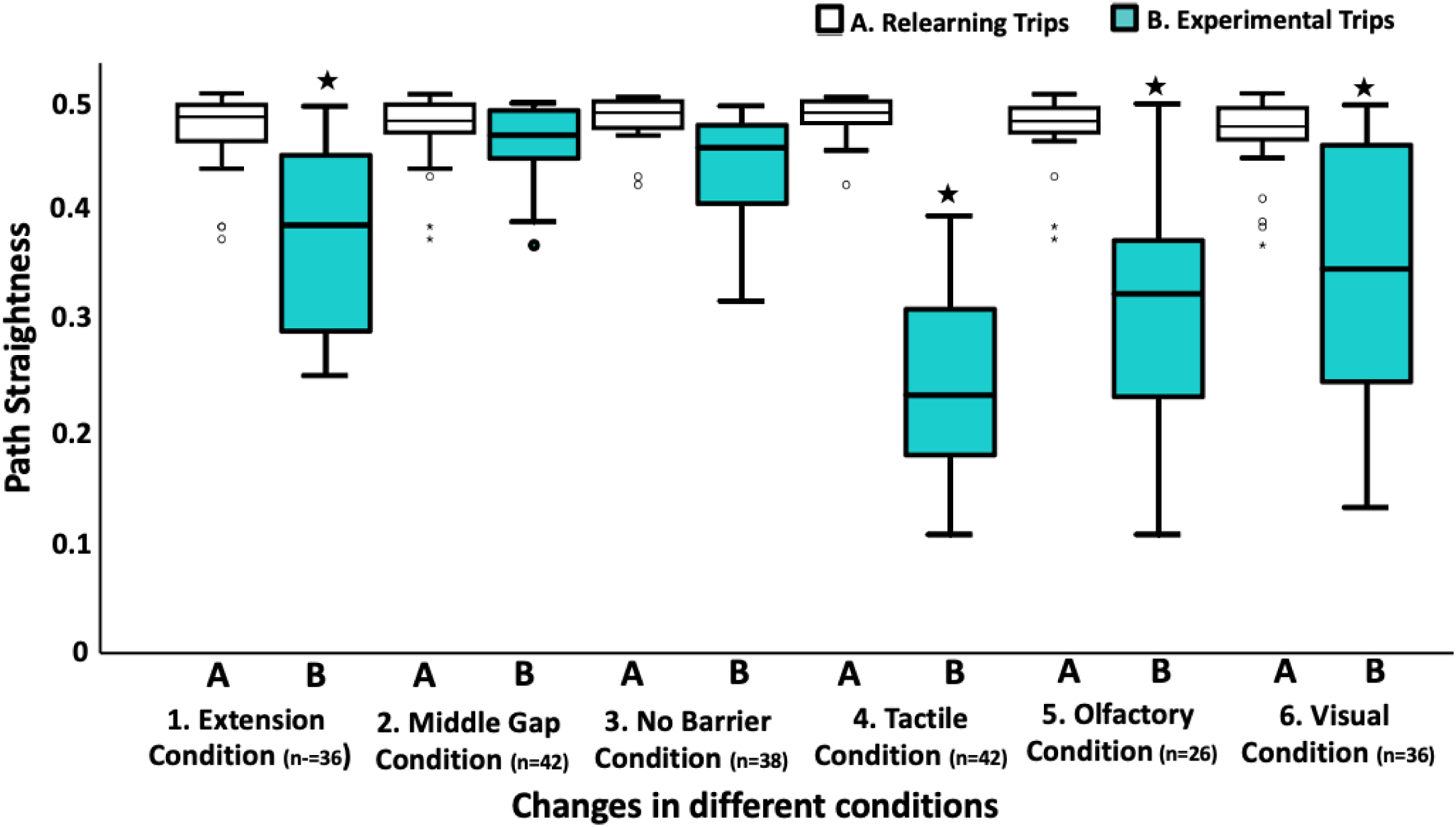
The path straightness of the nocturnal bull ants in different experimental conditions. The ‘*’ indicates that the path straightness of a condition was significantly different from the relearning trips of the same condition.

**Fig. 8.**
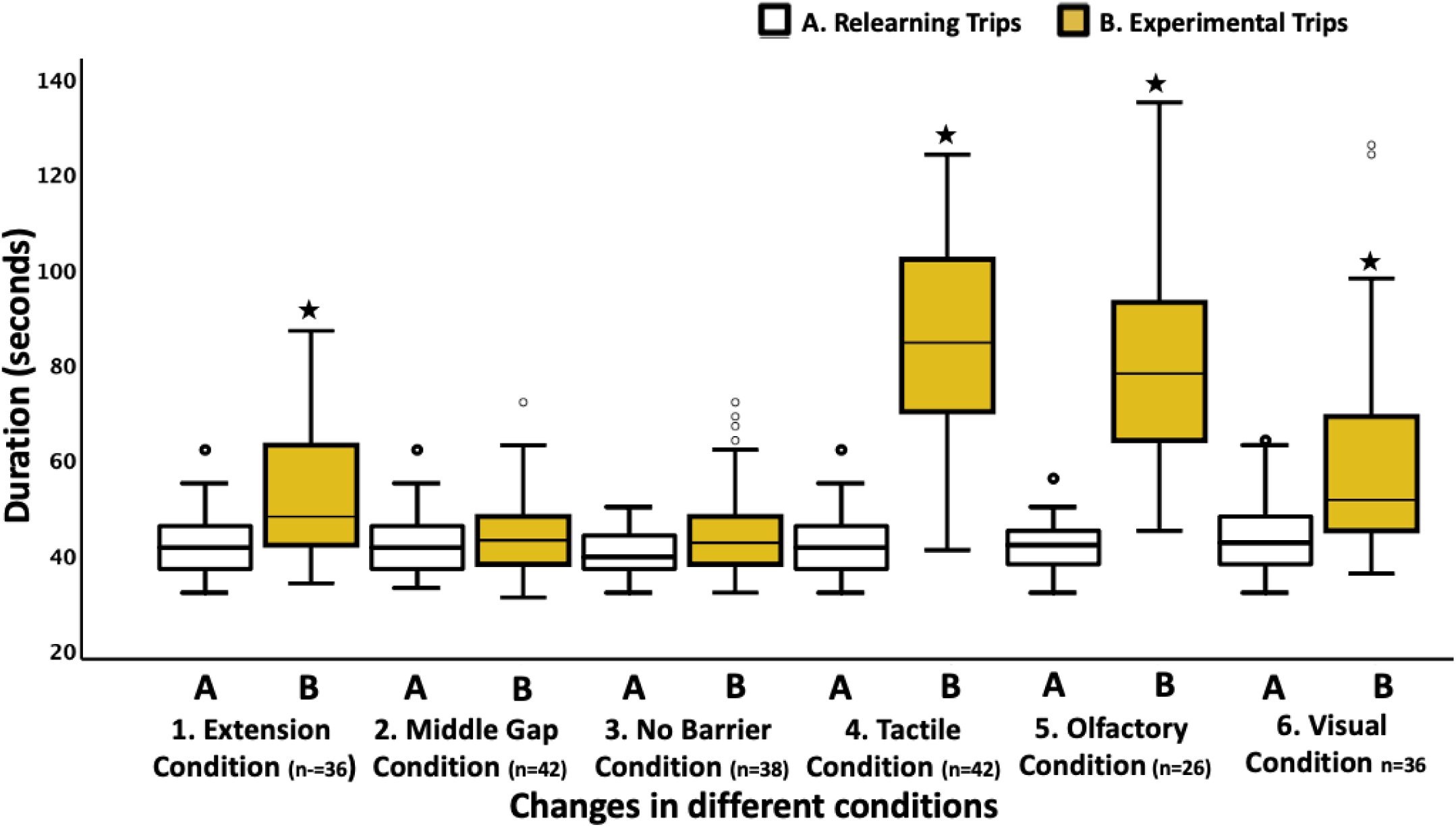
The time taken to reach 1.5 m above the nest entrance of the nocturnal bull ants in different experimental conditions. The ‘*’ indicates that the duration of a condition was significantly different compared to the relearning trips of the same condition.

#### Effect of tactile changes on learning

When foragers encountered the tactile changes on their foraging tree trunk, they mostly avoided the added cloth and moved around it (Fig. 6D). The ants stopped, scanned, and searched for an alternative way to reach the tree canopy (figure S3). As a result, paths were less straight (paired t test, t (39) = 12.56, p < 0.001) and of longer duration (paired t test, t (39) = –3.468, p < 0.001) compared to the last relearning trip before the Tactile Condition (Fig. 7 & Fig. 8).

#### Effect of olfactory changes on learning

In the Olfactory Condition, individual foragers similarly avoided the new odour and attempted to move around it (Fig. 6E). Foragers stopped just before the first odour line, meandered around the odour line, and then ascended the tree. As a result, trips were less straight (paired t test, t (25) = 9.05, p < 0.001) and took more time (paired t test, t (25) = – 7.146, p < 0.001) compared to the relearning trips (Fig. 7 and 8).

#### Effect of visual changes on learning

Foragers were also affected in the Visual Condition. Paths appeared similar to the relearning trials until the ants approached the cloths on the sides of the tree (figure 6F). Hesitancy, meandering, and scanning then set in (Fig. 7 & Fig. 8). As a result, foragers were less straight (paired t test, t (35) = 4.306, p < 0.001) and took more time (paired t test, t (35) = –4.83, p = 0.001) compared to the relearning trips. After passing halfway from the bottom of the tree trunk to the barrier, foragers observed the visual changes. Most of the foragers stopped, scanned, and then moved around the barrier to reach the canopy (Fig. 6 and Fig. S3).

## DISCUSSION

Our findings demonstrate both how nocturnal bull ants learn to avoid obstacles in their environment and how they adapt their learned motor routine to novel sensory stimuli. When the foragers confronted the barrier on their foraging corridor for the first time, almost half of them failed to overcome the barrier. In subsequent trips, the foragers became more successful. In subsequent detour trials, foragers became faster to pass the barrier and paths became straighter. Some foragers modified their foraging paths quickly, even within two trials, while other ants changed the paths more gradually over trips. When the position of the barrier was changed or removed altogether, foragers predominantly showed their learned motor routine, detouring with a similar path as before. We also found that when foragers detected the tactile or olfactory changes, they immediately showed neophobic behaviour, scanning and avoiding those novel cues. Visual changes on their learned route also adversely affected their navigation and motor routine.

The tested ants improved from 54% success on their first detour trial to 98% success on their 8^th^. Studies on other animals have shown a similar progression in increased success rates in multiple trials of detouring around a barrier on their way to a goal. In horses (*Equus caballus*), domestic dogs (*Canis familiaris),* domestic chicks, mice and dingoes, more than 50% individuals were successful in their first attempt and the success rate improved over trials (Pongrácz et al., 2001; Pongrácz et al., 2005; Baragli et al., 2011, Juszczak et al., 2016). Many studies found improved detouring over trials (Pongrácz et al., 2001; Wallis et al. 2001; Parker et al., 2005; Smith and Litchfield, 2010; Boogert et al., 2011; Anderson et al., 2016). In our detour learning experiments, foragers became faster to pass the barrier and walked straighter paths over several trials. Most of them veered left or right on the trunk well before the barrier. Several studies have found that animals become faster in reaching the destination around a barrier over several trials (Spigel, 1964; Vallortigara et al., 1988; Lockman et al., 2001; Parker et al., 2005; Smith and Litchfield, 2010; Baragli et al., 2011; Juszczak et al., 2016).

Our experiments found that ants maintained their newly learned route when the position of the barrier was changed, or the barrier was removed altogether. The ants started at a similar position at the bottom of the tree before and after the barrier was put in place, although with the barrier, most ants headed to a different position on the tree, one edge of the barrier. Retaining the learned new route suggests that foragers were not solely using cues of the barrier to navigate around it. They could have used a combination of a local vector [67] from the bottom of the tree to the edge of the barrier and using surrounding visual cues for route guidance. Two desert ants, *Melophorus bagoti* and *Cataglyphis fortis*, also maintain their newly learned route, which many ants learned after they encountered a pit trap on the direct route home (Collett et al., 1998). The trap was not visible until the ants were right at the edge, so that on the first run with the trap, homing ants fell into the trap. Ants that learned to detour around the trap persisted in their newly learned routes even after the pit was removed. Reinforcement learning was posited to play a key role in maintaining such learned routes (Collett et al., 1998) and reinforcement learning likely played a role in maintaining the new routes of the bull ants in this study as well.

When the nocturnal bull ants encountered olfactory and tactile changes on the tree trunk, the navigation of the experienced foragers was profoundly affected. Ants avoided the changed terrain. We interpret such behaviours tentatively as neophobia, an interpretation that requires further study. Novel objects are commonly used to assess neophobia, generally by placing an object near a familiar food source. Fear of novel environments (Mettke-Hofmann et al., 2009; Cohen et al., 2015; Elvidge et al., 2016) and fear of novel objects (Greenberg, 1988; Mettke-Hofmann et al., 2002; Moretti et al., 2015; Brown et al., 2016) produce a conflict between the motivation to feed and the motivation to avoid the object. Trained desert ants, *Cataglyphis fortis,* use odours from a food source or at the nest in navigation and searching and use the olfactory cues as route guidance in the absence of other navigational information (Wolf and Wehner, 2000; Buehlmann et al. 2015). Tactile learning experiments on *C. fortis* showed that the ants use tactile cues of the ground structure (Seidl and Wehner, 2006). With a novel or altered ground mark, however, these ants exhibited avoidance responses, likewise suggesting neophobia.

As discussed, the tested ants were affected adversely when a part of the surrounding visual scene was blocked. Changes of visual panorama may affect the navigation of ants, sometimes severely, although their performance recovers within a few days or over several trials (Graham and Cheng, 2009; Wystrach et al., 2011; Julle-Daniere et al., 2014; Narendra and Ramirez-esquivel, 2017; Freas et al., 2018a; Freas and Cheng, 2019; Islam et al., 2020). Even small visual changes such as the removal of three trees in a forest of trees may adversely affect bull ants (Narendra and Ramirez-esquivel, 2017). The ants did not require the surrounding visual cues to the sides of the tree as they still managed to negotiate the barrier with those cues blocked, albeit with more meandering and scanning. Meandering and scanning increase when ants encounter visual changes (Narendra and Ramirez-esquivel, 2017; Islam et al., 2020) and are interpreted as behaviours for learning the changed cues. Thus, we do not interpret the ants’ reactions as neophobia in this case.

Taken together, our results suggest that individual foragers learn a new route around a barrier on the tree, gradually in most cases, which is indicative of spatial learning. Most of the ants did not rely on barrier-based cues but rather used a local vector and probably surrounding visual cues to detour. Our findings also suggest a large initial neophobic response to tactile and olfactory changes on the route.

## Authors Contribution

MI, KC: conceptual development; MI, JFK, KC: experimental design; MI, SD: data collection; MI, ZM, SD: data analysis; MI, KC: original manuscript; MI, SD, ZM, JFK, KC: review & editing; all authors gave final approval for publication.

## Acknowledgements

We thank Macquarie University for giving us access and support in the field site on campus and are grateful to the Neuroethology Lab Group for valuable discussions in lab meetings.

